# Omicron and Alpha P680H block SARS-CoV2 spike protein from accessing cholinergic inflammatory pathway via α9-nAChR mitigating the risk of MIS-C

**DOI:** 10.1101/2022.02.18.481096

**Authors:** Ulises Santiago, Carlos J. Camacho

## Abstract

Sequence homology between neurotoxins and the site encompassing the furin cleavage site _680_SPRRAR_685_ in the spike protein (S) of CoV2 suggested that this site could interact with nicotinic acetylcholine receptors (nAChRs). Molecular dynamics simulations confirm robust structural similarity between wild-type (WT) CoV2 and the binding motif of α-conotoxin to α9 nAChR, which is known to modulate IL-1β in immune cells. We show that the structural integrity of this binding motif is eliminated by Alpha P681H mutation, reemerged in Delta variant P681R, and disappeared again with Omicron N679H/P681H. Interactions between the toxin-mimic CoV2 motif and α9-nAChR are expected to trigger the release of pro-inflammatory cytokines an effect that is mollified by Alpha and Omicron. Clinical features of this interaction site are relevant because, contrary to most regions in the S protein, the furin binding site does not appear to trigger an immune response prior to cleavage, indicating that the cholinergic pathway should be activated in the respiratory tract and nasal mucosa where α9-nAChR co-localizes with the virus. The correlation of changes on this motif by the different variants closely matches the reported cases of Multisystem Inflammatory Syndrome in Children by the CDC, and predicts significant mitigation of MIS-C with the Omicron variant. Our findings strongly motivate further study of this cholinergic pathway as one source of the cytokine storm triggered by CoV2.

---

Three years into the COVID-19 pandemic has brought little mechanistic understanding of the inflammatory pathways and cytokine storm triggered by the virus^1,2^. However, the relatively rapid appearance of new variants of SARS-CoV2 offers clues to how and why it is adapting^3^. Significant efforts are being devoted to understanding what makes the virus more infectious or transmissible^4^, and how it evolves to overcome a host’s immune response^5^. In the conventional paradigm of humoral immune responses, B cells recognize exposed conformational epitopes of protein antigens through interactions with surface expressed immunoglobulin receptors^6^. Based on sound thermodynamic principles, the stability of key structural motifs^7^ in these epitopes, and their availability^8^, or exposure, to bind are essential properties to understand the diverse humoral immune responses induced by relevant three-dimensional epitopes, as well as host-pathogen interactions^9^.

Significant among the multiple CoV2 mutations that have been fixated in the population is a unique proline located at P681 of the Spike (S) protein, which not only belongs to the most immunogenic of the epitopes of 2019 coronavirus disease patients^10,11^ that has been mutated in new variants but also forms part of the essential furin cleavage site _680_*S**P**RRAR*←S1/S2→*SVAS*_689_^12,13^. Unique to CoV2, this site recruits furin for cleavage of S proteins into S1/S2, a process that is critical for the subsequent cleavage by TMPRSS2 at site S2’ to trigger cell fusion and viral entry^4,14^. Variants of concern (VOC) of CoV2 have mutated this amino acid twice: Alpha and Omicron (P681H) and Delta (P681R). The site is in a flexible loop of S^15^, making the proline mutation viable. Intriguingly, high immunogenicity at this epitope has only been observed at the C-end terminal of the S1, while no antigenicity has yet been reported neither for the intact nor for the N-terminal end of S2^10,11^.

Independently, based in part on reports of low smoking prevalence among hospitalized COVID-19 patients^16^, the well-established role of nicotinic acetylcholine receptors (nAChRs) in inflamation^17^, and a distant sequence similarity between a neurotoxin motif that targets nAChRs and a region of the S protein of SARS-CoV2 near the furin cleavage site (**Fig. 1A**)^18^, increasing attention has been paid to the hypothesis that nAChRs might have some involvement in the clinical manifestations of COVID-19^16,19–21^. One of them being systemic hyper inflammatory syndrome^22^. The expression of α7-nAChR subunit in the nervous and immune system has suggested this subunit as a possible target^19,23^. Much less attention has been given to the α9-nAChR subunit.

**Figure 1.**
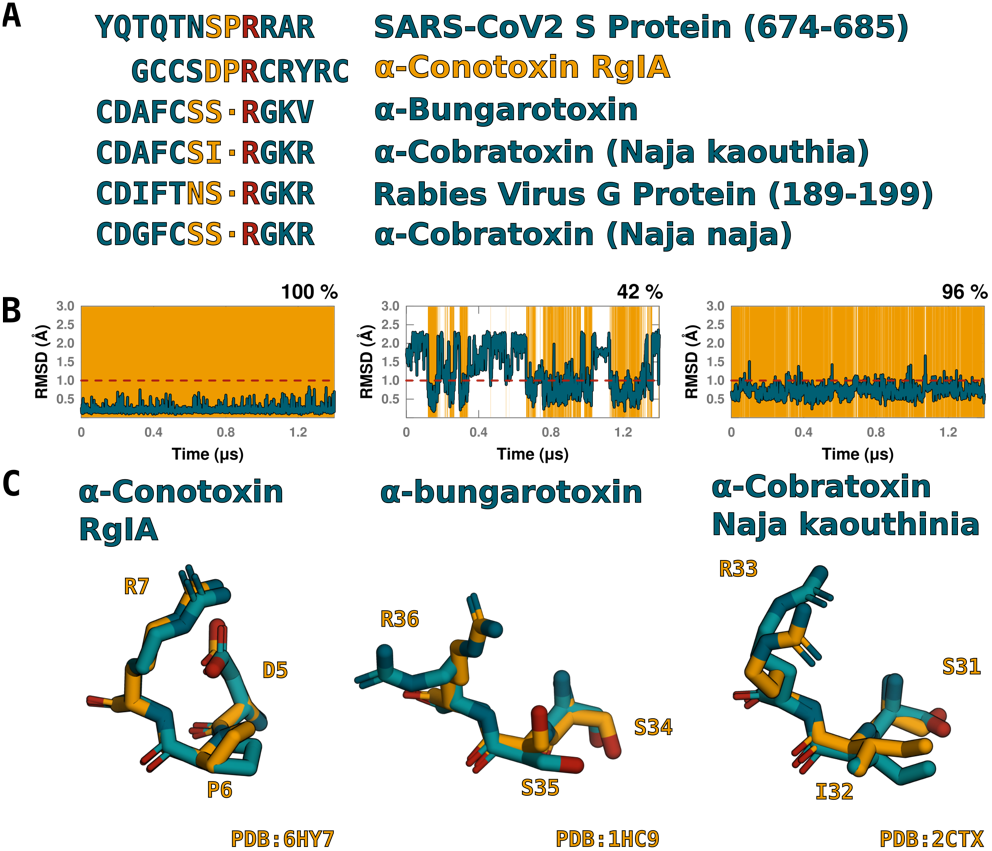
Sequence alignment of furin binding site of SARS-CoV2 and structural stability analysis of toxins that interact with nAChRs. **(A)** Besides previously noted toxins, we discovered α-conotoxin that shares the critical PR motif with CoV2. **(B)** MD simulations of 4 concatenated 350 ns production runs (see Methods) show that, in the absence of the receptor, toxins are stable and all have a critical interacting arginine homolog to R682 of CoV2. **(C)** Most stable cluster center closely mimics bound structure of three toxins with known structure.

The α9-nAChR subunit is found in the pituitary, tongue, olfactory epithelium, and hair cells of the cochlea^17^. Hence, in a more direct path of contact with the CoV2 virus. Additionally, it has been reported in a variety of native immune cells, including B cells^24^, and T lymphocytes^25^, strongly suggesting that they are involved in immunological processes in the respiratory airways. More importantly, α9 nAChRs have been shown to modulate the release of IL-1β^26–28^.

In order to gain molecular insights on the possible connection of SARS-CoV2 and nAChRs, we studied a panel of co-crystal structures with the neurotoxin binding motif. We found that all the motifs entail a conserved Arginine anchor residue^7^. Molecular dynamics (MD) of the corresponding sequences confirm that this Arginine is bound-like and exposed to solvent prior to binding, consistent with the expected properties of the motif responsible for molecular recognition^7^. Strikingly, we found similar structural properties in the intact (pre-cleavage) loop around the conserved R682 in the furin binding site of the S protein of wild-type (WT) SARS-Cov2. Namely, the motif _680_SPR_682_ of S perfectly matches the DPR interacting sites of α-conotoxin bound to α9-nAChR subunit (PDB 6HY7)^29^. Interestingly, the bound-like motif of R682 is lost after cleavage. Thus, the predicted interaction between S and α9 nAChR should only happen prior to cleavage, i.e., when the virus resides in the respiratory track and nasal mucosa. Of note, this host-pathogen interaction in the furin binding site should prevent cleavage, slowing down cell fusion and viral entry^4,14^. MDs of the mutated Alpha variant P681H eliminates the structural integrity of the α-conotoxin binding motif by inducing a cation-pi stacking interaction between R681 and H681, which no longer fits in the α9 nAChR binding pocket. In contrast, the Delta variant P681R resuscitates the α-conotoxin binding motif, but the increased flexibility of R681 relative to P681 makes the motif less stable than WT. Finally, Omicron N679K/P681H recapitulates the stacking of H681 and R682, again preventing an interaction with α9 nAChR. While generic nAChR agonists like nicotine, or acetylcholine inhibit IL-1β release, and reduce inflammation^17^, α9-nAChR antagonists, such as α-conotoxin or the toxin-mimic spike protein of WT, beta, gamma, iota, and to a lesser extent the Delta variant of CoV2, should trigger the release of pro-inflammatory cytokines. Strikingly, our findings show that both Alpha and Omicron variants block this cholinergic inflammatory pathway. These predictions match closely CDC reported data of Multisystem Inflammatory Syndrome in Children (MIS-C). Namely, MIS-C drastically diminished during the Alpha variant; a mild wave of cases reappeared when the variant of concern (VOC) was Delta; and, here, we predict that this pathway to inflammation will be fully blocked with Omicron significantly diminishing the risk of MIS-C.

## RESULTS

### Sequence and structural homology of neurotoxins and furin binding site

As shown by the panel of co-crystal structures in **Fig. 1B**, the neurotoxin binding motifs entail a conserved Arginine extending away from a loop, which is known to interact with nAChRs. The toxins are stabilized by disulfide bonds, giving rise to relatively stable and potent binding motifs. As expected, MDs show that, even before encountering its receptor^30^, the toxin motifs acquired their bound-like conformation (**Fig. 1C**). Of particular interest is the highly stable α-conotoxin that binds α9 nAChR subunit^29^ (**Fig. 2A**) because the homology of DPR with _680_SPR_682_ in the spike protein of CoV2 (**Fig. 1**).

**Figure 2.**
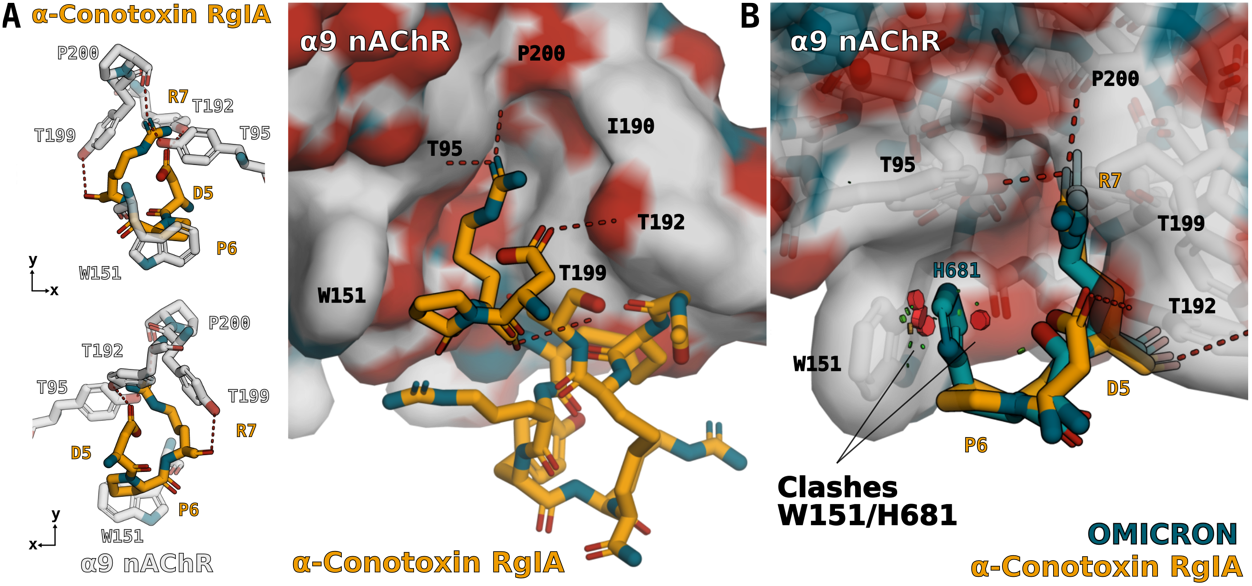
Alpha and Omicron _680_SHR_682_ motif does not fit in α9-nAChR binding pocket. **(A)** Co-crystal of α-conotoxin and α9-nAChR subunit (PDB 6HY7). **(B)** Stacking of H681 and R682 overlapped with co-crystal shows clashes (red cylinders) between spike H681 and α9-nAChR subunit W151.

**Figure 3A** shows the root-mean-square-deviations (RMSD) between MD snapshots of 6 independent concatenated runs totaling more than 2 μs of the DPR binding motif of α-conotoxin and _680_SPR_682_, _680_SHR_682_, _680_SRR_682_, and (K)_680_SHR_682_ motifs of WT, Alpha, Delta and Omicron variants, respectively. Although there are no disulfide bonds restraints in the spike furin loop, the WT structure consistently samples the toxin-like α9 nAChR binding motif (**Fig. 3B**), displaying a remarkable structural and chemical similarity with all the interacting atoms observed in co-crystal (**Fig. 2A**). Note that the RMSD is almost as accurate as the MDs of the toxin itself in **Fig. 1BC**. This observation strongly suggests that, like α-conotoxin, the spike protein should readily bind the α9-nAChR subunit. Naturally, the same applies to all the sequences that don’t mutate the furin binding site, i.e., beta, gamma, iota and other variants.

**Figure 3.**
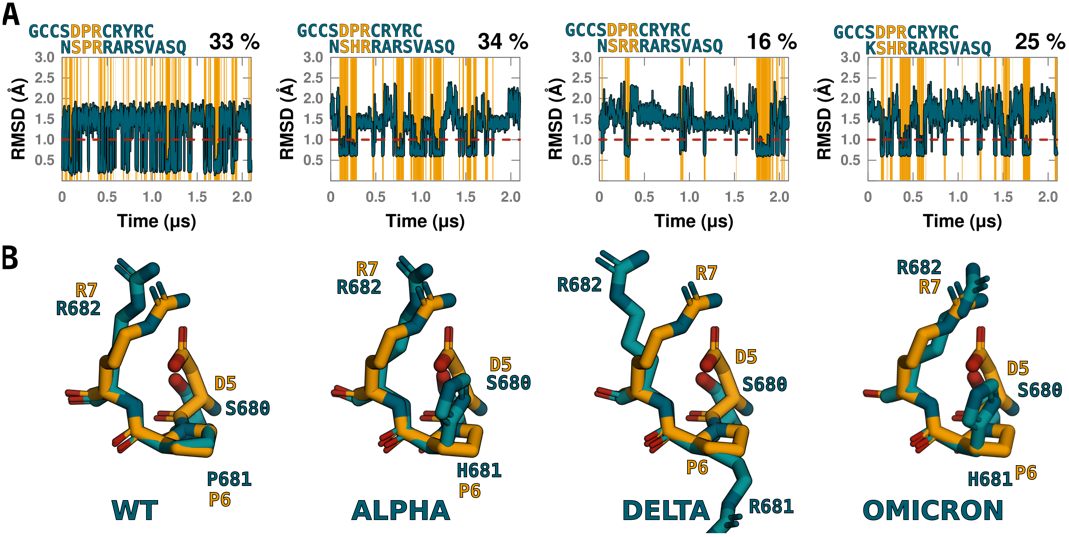
CoV2 variants mimic α-conotoxin binding motif to α9-nAChR. **(A)** RMSD of backbone and C_β_ of toxin DPR bound structure relative to WT SPR and VOC SHR and SRR of SARS-CoV2 for 6 independent MDs of 350 ns each (see Methods). **(B)** Cluster of snapshots below 1 Å RMSD show a binding motif similar to α-conotoxin, but Alpha and Omicron histidine is stacked parallel to R682.

### Alpha and Omicron variant P681H no longer fits in α9 nAChR binding pocket

As already mentioned P681 has been mutated by VOC Alpha, Delta and Omicron. A natural question to ask is: what is the impact of these mutations in the α9-nAChR binding motif? To answer this question, we repeated the stability analysis performed for WT, and surprisingly found that both Alpha and Omicron are able to acquire an overall similar structural motif as α-conotoxin (**Fig. 3AB**). Of note, the dynamics of the binding motif is different than WT, i.e., the bound-like state is stable for somewhat longer time scales. As shown in **Fig. 3B**, this extra stability arises due to a cation-pi interaction betwen the histidine aromatic ring and R682. As shown in **Fig. 2B**, this motif does not fit in the pocket between W151 and Y199. We note that W151 is an essential motif of the α9-nAChR binding pocket, and, in general, this side chain has little mobility. Hence, we conclude that neither the Alpha nor the Omicron variant are able to bind the α9 nACh receptor.

### The α9-nAChR binding motif reemerges with the Delta mutation P681R

Though less dominant due to extra flexibility surrounding R681, Delta acquires the toxin-like motif for about 16% of the simulation time (**Fig. 2A**). The most common structural motif, with a RMSD of about 1.3 Å, differs from the toxin motif shown in **Fig. 2B** on S680 being flipped outwards, which could potentially rearrange after R682 binds in the pocket. Overall, the expectation is that Delta should still be able to engage α9-nAChR, though less effectively than WT.

## DISCUSSION

Three years into the COVID-19 pandemic has brought little mechanistic understanding of the inflammatory pathways and cytokine storm triggered by the virus^1,2^. Here, we provide strong evidence that the physical interaction of the furin binding site of the spike protein of SARS-CoV2 and the α9 subunit of the nicotinic acetylcholine receptor (α9-nAChR) should activate the cholinergic inflammatory pathway.

More importantly, α9-nAChR is known to modulate the release of IL-1β, as well as induced (mucosal) IL-6^31^ and other pro-inflammatory cytokines. Transcripts of α9-nAChR have been shown to co-localize^17^ with the access point of the virus. Therefore, contrary to other nAChRs, the α9 subunit meets the virus in the respiratory track and nasal mucosa.

Remarkable, all variants that do not mutate the region around the furin cleavage site _680_S**P**R_682_ have a strong sequence and structural similarity with a potent toxin motif (**Fig. 3AB**) that should readily recognize and bind α9-nAChR. Interactions between the α-conotoxin-mimic CoV2 motif and α9 nAChR are expected to trigger the release of pro-inflammatory cytokines. This interaction at the furin binding motif should also slow down the cleavage by furin. This is important since it is known that that furin cleavage occurs even during virus packaging^4^. And, after cleavage, the C-end terminal of S1 that encompass _680_S**P**R_682_ motif is not only highly immunogenic, but also its structure is no longer compatible a toxin-binding motif capable of binding α9 nAChR (**Fig. S1**).

The P681H mutation of the Alpha variant keeps a motif similar to α-conotoxin, with the important caveat that the aromatic ring of H681 forms a stacking interaction with R682. This motif no longer fits in the α9-nAChR binding pocket (**Fig. 2B**), preventing the interaction between the Alpha variant and the cholinergic inflammatory pathway via the α9 subunit. The same behavior is observed for Omicron, whereas Delta partially recovers the toxin binding motif, and, thus, some capacity to trigger this inflammatory pathway. As shown in **Fig. 4**, our findings accurately track the appearance of these different variants of concern and reported cases of Multisystem Inflammatory Syndrome in Children by the CDC. Our findings predict that Omicron has a significantly reduced risk to trigger MIS-C. Of note, the Alpha variant already showed a diminished risk of MIS-C. However, as noted in **Fig. 4**, the alpha variant always has some overlap with beta, gamma, iota and mu variants that are predicted to engage the cholinergic inflammatory pathway the same as wild-type SARS-CoV2.

**Figure 4.**
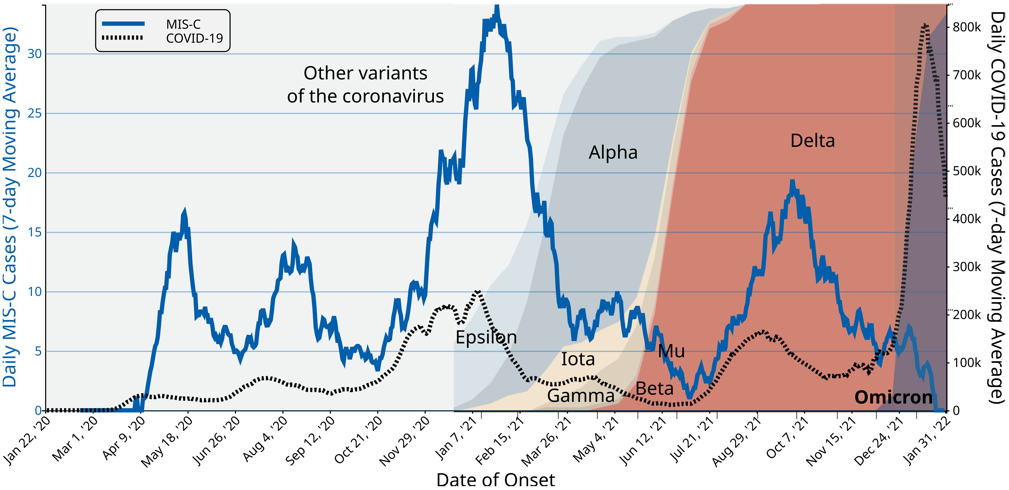
Daily MIS-C and COVID-19 cases as reported by the CDC correlated with variants of concern (from NYT tracking). The most significant feature is the significant drop of MIS-C when the VOC was Alpha. Note that Alpha never fully overcame other variants such as beta, gamma, etc. that can engage α9-nAChR. MIS-C had a weaker response to Delta. Only Alpha, Delta and Omicron had mutated the furin binding site. We predict that there will be no delayed response of MIS-C to Omicron.

Collectively, our findings validate molecular modeling as an approach to probe the host/pathogen interactions. This advance allows us to test the functional impact of past and future mutations by *in silico* scanning physical interactions. Based on the insights borne out by the structural evolution at the furin cleavage site, we surmise that Omicron eliminated the risk of MIS-C. The ultimate question of whether the N679K/P681H mutations efficiently evade an immune system response, have an unforeseen functional selective pressure, or benefit from simply keeping naive populations available for replication remains to be determined.

## METHODS

The initial peptide structures were generated in Pymol. The N- and C-termini were capped with ACE and NHE, respectively, except in peptides of the C-end terminal of S1 that were capped with ACE and -O. The molecular dynamics (MD) simulations were run with pmemd.cuda^32–34^ from AMBER18 using AMBER ff14SB force field^35^. We used tLeap binary (part of AMBER18) for solvating the structures in an octahedral TIP3P water box with a 12 Å distance from structure surface to the box edges, and closeness parameter of 0.75 Å. The system was neutralized and solvated in a solution of 150 mM NaCl. Simulations were carried out equilibrating the system for 1 ns at NPT using Berstein barostat to keep constant pressure at 1 atm at 300K, followed by 500 ns NPT production at 300 K, with non-bonded interaction cut off at 10 Å. Hydrogen bonds were constrained using SHAKE algorithm and integration time-step at 2 fs. Analysis of our repeated MDs, indicated that 150 ns was an adequate time for equilibration, and therefore the first 150 ns of all our runs were discarded from the analysis. Clustering and RMSD calculations were generated using Cpptraj^36^ software. Clusters were calculated considering the largest number of elements around a centroid with a distance less than a reported value (see text).

## ACKNOWLEDGEMENT

We are grateful for motivating and insightful discussions with Professors Maria Chikina, Dana Ascherman, and Wesley Pedgen. Comments to the MS by Gaby Gerlach and Alex Hammels are acknowledged.

## References

1. Yang, L. et al. The signal pathways and treatment of cytokine storm in COVID-19. Signal Transduct Target Ther 6, 255 (2021).

2. Fara, A., Mitrev, Z., Rosalia, R.A. & Assas, B.M. Cytokine storm and COVID-19: a chronicle of pro-inflammatory cytokines. Open Biol 10, 200160 (2020).

3. Callaway, E. Beyond Omicron: what’s next for COVID’s viral evolution. Nature 600, 204–207 (2021).

4. Peacock, T.P. et al. The furin cleavage site in the SARS-CoV-2 spike protein is required for transmission in ferrets. Nat Microbiol 6, 899–909 (2021).

5. Eguia, R.T. et al. A human coronavirus evolves antigenically to escape antibody immunity. PLoS Pathog 17, e1009453 (2021).

6. Benjamin, D.C. et al. The antigenic structure of proteins: a reappraisal. Annu Rev Immunol 2, 67–101 (1984).

7. Rajamani, D., Thiel, S., Vajda, S. & Camacho, C.J. Anchor residues in protein-protein interactions. Proc Natl Acad Sci U S A 101, 11287–92 (2004).

8. Travers, T.S. et al. Extensive Citrullination Promotes Immunogenicity of HSP90 through Protein Unfolding and Exposure of Cryptic Epitopes. J Immunol 197, 1926–36 (2016).

9. Camacho, C.J., Katsumata, Y. & Ascherman, D.P. Structural and thermodynamic approach to peptide immunogenicity. PLoS Comput Biol 4, e1000231 (2008).

10. Li, Y. et al. Linear epitope landscape of the SARS-CoV-2 Spike protein constructed from 1,051 COVID-19 patients. Cell Rep 34, 108915 (2021).

11. Li, Y. et al. Linear epitopes of SARS-CoV-2 spike protein elicit neutralizing antibodies in COVID-19 patients. Cell Mol Immunol 17, 1095–1097 (2020).

12. Hoffmann, M., Kleine-Weber, H. & Pohlmann, S. A Multibasic Cleavage Site in the Spike Protein of SARS-CoV-2 Is Essential for Infection of Human Lung Cells. Mol Cell 78, 779–784 e5 (2020).

13. Xing, Y., Li, X., Gao, X. & Dong, Q. Natural Polymorphisms Are Present in the Furin Cleavage Site of the SARS-CoV-2 Spike Glycoprotein. Front Genet 11, 783 (2020).

14. Bestle, D. et al. TMPRSS2 and furin are both essential for proteolytic activation of SARS-CoV-2 in human airway cells. Life Sci Alliance 3(2020).

15. Walls, A.C. et al. Structure, Function, and Antigenicity of the SARS-CoV-2 Spike Glycoprotein. Cell 181, 281–292 e6 (2020).

16. Farsalinos, K. et al. Editorial: Nicotine and SARS-CoV-2: COVID-19 may be a disease of the nicotinic cholinergic system. Toxicol Rep 7, 658–663 (2020).

17. Hone, A.J. & McIntosh, J.M. Nicotinic acetylcholine receptors in neuropathic and inflammatory pain. FEBS Lett 592, 1045–1062 (2018).

18. Changeux, J.P., Amoura, Z., Rey, F.A. & Miyara, M. A nicotinic hypothesis for Covid-19 with preventive and therapeutic implications. C R Biol 343, 33–39 (2020).

19. Oliveira, A.S.F. et al. A potential interaction between the SARS-CoV-2 spike protein and nicotinic acetylcholine receptors. Biophys J 120, 983–993 (2021).

20. Cheng, M.H. et al. Superantigenic character of an insert unique to SARS-CoV-2 spike supported by skewed TCR repertoire in patients with hyperinflammation. Proc Natl Acad Sci U S A 117, 25254–25262 (2020).

21. Cheng, M.H. et al. A monoclonal antibody against staphylococcal enterotoxin B superantigen inhibits SARS-CoV-2 entry in vitro. Structure 29, 951–962 e3 (2021).

22. Gruber, C.N. et al. Mapping Systemic Inflammation and Antibody Responses in Multisystem Inflammatory Syndrome in Children (MIS-C). Cell 183, 982–995 e14 (2020).

23. Courties, A. et al. Regulation of the acetylcholine/alpha7nAChR anti-inflammatory pathway in COVID-19 patients. Sci Rep 11, 11886 (2021).

24. St-Pierre, S. et al. Nicotinic Acetylcholine Receptors Modulate Bone Marrow-Derived Pro-Inflammatory Monocyte Production and Survival. PLoS One 11, e0150230 (2016).

25. Peng, H. et al. Characterization of the human nicotinic acetylcholine receptor subunit alpha (alpha) 9 (CHRNA9) and alpha (alpha) 10 (CHRNA10) in lymphocytes. Life Sci 76, 263–80 (2004).

26. Hecker, A. et al. Phosphocholine-Modified Macromolecules and Canonical Nicotinic Agonists Inhibit ATP-Induced IL-1beta Release. J Immunol 195, 2325–34 (2015).

27. Richter, K. et al. Phosphocholine - an agonist of metabotropic but not of ionotropic functions of alpha9-containing nicotinic acetylcholine receptors. Sci Rep 6, 28660 (2016).

28. Zakrzewicz, A. et al. Canonical and Novel Non-Canonical Cholinergic Agonists Inhibit ATP-Induced Release of Monocytic Interleukin-1beta via Different Combinations of Nicotinic Acetylcholine Receptor Subunits alpha7, alpha9 and alpha10. Front Cell Neurosci 11, 189 (2017).

29. Zouridakis, M. et al. Crystal Structure of the Monomeric Extracellular Domain of alpha9 Nicotinic Receptor Subunit in Complex With alpha-Conotoxin RgIA: Molecular Dynamics Insights Into RgIA Binding to alpha9alpha10 Nicotinic Receptors. Front Pharmacol 10, 474 (2019).

30. Kimura, S.R., Brower, R.C., Vajda, S. & Camacho, C.J. Dynamical view of the positions of key side chains in protein-protein recognition. Biophys J 80, 635–42 (2001).

31. Cahill, C.M. & Rogers, J.T. Interleukin (IL) 1beta induction of IL-6 is mediated by a novel phosphatidylinositol 3-kinase-dependent AKT/IkappaB kinase alpha pathway targeting activator protein-1. J Biol Chem 283, 25900–12 (2008).

32. Case, D.A. et al. The Amber biomolecular simulation programs. J Comput Chem 26, 1668–88 (2005).

33. Gotz, A.W. et al. Routine Microsecond Molecular Dynamics Simulations with AMBER on GPUs. 1. Generalized Born. J Chem Theory Comput 8, 1542–1555 (2012).

34. Salomon-Ferrer, R., Gotz, A.W., Poole, D., Le Grand, S. & Walker, R.C. Routine Microsecond Molecular Dynamics Simulations with AMBER on GPUs. 2. Explicit Solvent Particle Mesh Ewald. J Chem Theory Comput 9, 3878–88 (2013).

35. Maier, J.A. et al. ff14SB: Improving the Accuracy of Protein Side Chain and Backbone Parameters from ff99SB. J Chem Theory Comput 11, 3696–713 (2015).

36. Roe, D.R. & Cheatham, T.E., 3rd. PTRAJ and CPPTRAJ: Software for Processing and Analysis of Molecular Dynamics Trajectory Data. J Chem Theory Comput 9, 3084–95 (2013).

